# Multiplexed, bioorthogonal labeling of multicomponent, biomolecular complexes using genomically encoded, non-canonical amino acids

**DOI:** 10.1101/730465

**Authors:** Bijoy J. Desai, Ruben L. Gonzalez

**Affiliations:** Department of Chemistry, Columbia University, 3000 Broadway, MC3126, New York, NY 10027, USA

**Author notes:** To whom correspondence should be addressed: Ruben L. Gonzalez Jr., Department of Chemistry, Columbia University, 3000 Broadway, MC3126, New York, NY 10027, USA.

## Abstract

Stunning advances in the structural biology of multicomponent biomolecular complexes (MBCs) have ushered in an era of intense, structure-guided mechanistic and functional studies of these complexes. Nonetheless, existing methods to site-specifically conjugate MBCs with biochemical and biophysical labels are notoriously impracticable and/or significantly perturb MBC assembly and function. To overcome these limitations, we have developed a general, multiplexed method in which we genomically encode non-canonical amino acids (ncAAs) into multiple, structure-informed, individual sites within a target MBC; select for ncAA-containing MBC variants that assemble and function like the wildtype MBC; and site-specifically conjugate biochemical or biophysical labels to these ncAAs. As a proof-of-principle, we have used this method to generate unique single-molecule fluorescence resonance energy transfer (smFRET) signals reporting on ribosome structural dynamics that have thus far remained inaccessible to smFRET studies of translation.

---

Many fundamental cellular processes, including DNA repair, replication, transcription, messenger RNA (mRNA) processing and splicing, nuclear export, mRNA decay, translation, and protein degradation, are performed by MBCs. Continuing advances in X-ray crystallography and, more recently, cryogenic electron microscopy (cryo-EM) studies of MBCs are providing researchers with structural frameworks for most informatively positioning biochemical and biophysical probes and interpreting the resulting biochemical and biophysical data in terms of structure-based mechanistic models^1^. Unfortunately, however, it remains difficult and, in some cases, impossible to site-specifically label MBCs at defined positions, severely impeding mechanistic and functional studies. The primary reasons for this are that MBCs are composed of up to thousands of oftentimes essential components, have tightly controlled component stoichiometries, and are typically assembled through intricate and highly regulated assembly pathways^2–4^. Consequently, common methods for site-specifically labeling proteins, such as conjugation of a maleimide-derivatized reporter to a cysteine residue, are impracticable for MBCs that contain hundreds of native reactive residues^5^, while the peptide tags often used in chemo-enzymatic labeling methods are usually limited to protein termini where they are least likely to inhibit assembly and/or function^6, 7^. In addition, many labeling approaches involve production of a target protein from a genetic construct (*e.g.*, a plasmid) that removes the gene from its native genomic regulatory context, and this can perturb the component stoichiometry and/or cellular assembly process^8^. Attempts to overcome these issues by partially or fully *in vitro* reconstituting MBCs from recombinantly overexpressed, purified, and labeled components often results in compositionally and/or functionally heterogeneous MBC mixtures that exhibit impaired activities^8^.

Here, we report an approach that integrates homologous recombination-based multiplexed genome engineering (MGE)^9^, ncAA mutagenesis technology^10^, and bioorthogonal chemistry^11^ to rapidly generate numerous, fully functional MBC variants in which each variant can carry a biochemical or biophysical label at one or more defined target positions. Our approach combines the power of MGE to rapidly generate multiple, orthogonal codon mutations (Fig. 1a) with the specificity and modularity of bioorthogonal ncAA-based conjugation chemistry, while maintaining the genomic regulatory context, *in vivo* assembly pathway, and functional integrity of the target MBC (Fig. 1b-c). Because ncAA mutations are much less perturbative than peptide tag insertions and because small-molecule, chemistry-based conjugation is much less sterically restricted than enzyme-based conjugation, our approach allows efficient labeling of virtually any position in an MBC. Additionally, because our method requires that each strain carrying a variant MBC effectively competes with the strain carrying the wild-type MBC, our approach naturally counter-selects against variants that impair MBC function. Finally, the abundance of biochemical and biophysical probes that have been derivatized for bioorthogonal conjugation means that MBCs can be labeled with optical probes such as fluorophores and quenchers, affinity tags such as biotin and digoxigenin, radioisotopes, spin-labels, chemical probing agents, crosslinking agents, solid-state supports, nanoparticles, microspheres, *etc.*, for use in a nearly endless array of biochemical and biophysical applications (Fig. 1d).

**Figure 1.**
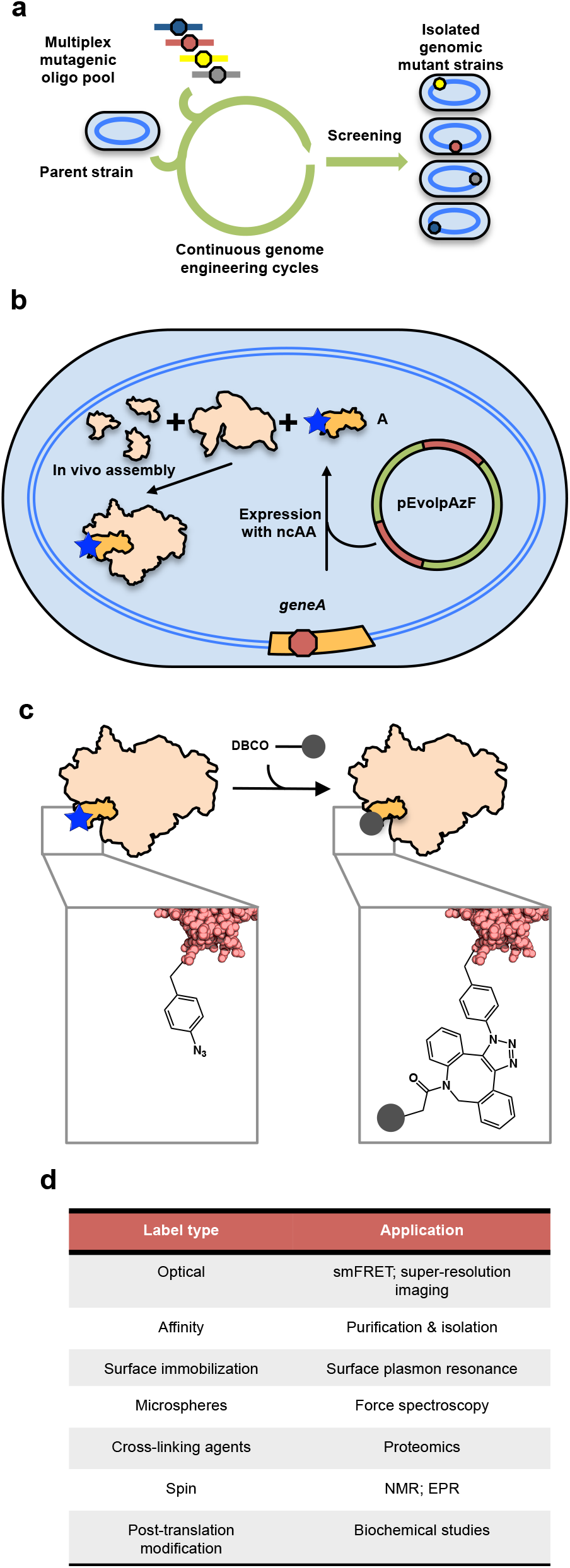
Using a combination of MGE, ncAA mutagenesis technology, and bioorthogonal chemistry to site-specifically label MBCs without perturbing their genomic regulation, *in vivo* assembly, structure, or function. (a) Iterative MGE cycles are used to introduce orthogonal codon mutations (hexagons) at specific genomic positions in genes encoding MBC proteins in a background of wildtype parent strain. (b) *In vivo* expression and assembly of the MBC in each mutant strain, including the MBC protein carrying the incorporated ncAA (blue star), is achieved by performing the MGE cycles in the presence of a plasmid expressing an ncAA-specific, orthogonal tRNA-tRNA synthetase pair and in the presence of the ncAA in the growth media such that each resulting mutant strain can assemble MBCs carrying the ncAA at one of the targeted positions. In our case, we have used *p*-AzF as the ncAA and the pEvol-pAzFRS.2.t1 plasmid to express the corresponding, orthogonal tRNA-tRNA synthetase pair. (c) ncAAs incorporated into MBCs purified from successfully selected mutant strains can be conjugated to an appropriately derivatized label or reporter (dark-grey) using bioorthogonal chemistry. In our case, we have used the strain-promoted, azide-alkyne, bioorthogonal conjugation reaction of *p*-AzF with DBCO-derivatized Cy3- and/or Cy5 fluorophores. (d) Examples of the different types of biochemical and biophysical labels and reporters that can be adapted to the approach described here and their respective applications in biochemical and biophysical studies of MBCs.

To demonstrate the power of our approach, we chose to target the *Escherichia coli* ribosome – a 2.5 MDa, two-subunit MBC comprised of 55 ribosomal proteins and 3 ribosomal RNAs (rRNAs). In bacteria, assembly and maturation of the ribosome is a complex process that requires the action of ~100 cellular factors^2^. The importance of the genomic regulation and *in vivo* assembly and maturation process of this MBC is highlighted by the observation that ribosomes composed of a fully *in vitro* reconstituted small, or 30S, ribosomal subunit are only 34% active compared to ribosomes composed of 30S subunits purified from cells^8^, a fact that has complicated interpretation of biophysical studies performed using ribosomes composed of 30S subunits that were site-specifically fluorophore-labeled *via* full *in vitro* reconstitution^12^. For site-specific labeling of the ribosome, we targeted thirteen different positions across nine ribosomal protein genes (Table 1, Supplementary Fig. 1), such that labeling with fluorophores at those positions would enable smFRET studies of three ribosomal structural rearrangements that have either been studied using smFRET signals that could not be interpreted unambiguously^13, 14^ or that have otherwise remained inaccessible to smFRET studies: (*i*) intra-subunit rotation of the ‘head’ domain of the 30S subunit relative to the ‘body’ domain (*i.e.*, ‘head swiveling’, HS), (*ii*) movement of a translating ribosome along its mRNA template (*i.e.*, ‘mRNA translocation’, MT), and (*iii*) rotation of the large, or 50S, ribosomal subunit relative to the 30S subunit (*i.e.*, ‘intersubunit rotation’, IR) (Supplementary Fig.1).

**Table 1.** Percent enrichment of mutants after MGE

| smFRET signal | Target for mutation | Number of bp mutated | Number of MGE cycles | Percent enrichment<sup>1</sup> |
| --- | --- | --- | --- | --- |
| HS1 | 30S S7G112 | 3 | 8 | 1.0 |
|  | 30S S11A102 | 3 | 6 | 1.0 |
| HS2 | 30S S7K131 | 2 | 8 | 2.1 |
|  | 30S S18R8 | 2 | 6 | 2.1 |
| HS3 | 30S S7G112 | 3 | 8 | 1.0 |
|  | 30S S18Q75 | 1 | 8 | 0 |
| HS4 | 30S S19Q56 | 1 | 8 | 3.1 |
|  | 30S S12K108 | 2 | 6 | 4.6 |
| HS5 | 30S S13D11 | 2 | 6 | 0 |
|  | 30S S11R106 | 3 | 8 | 0 |
| MT1 | 30S S18R8 | 2 | 6 | 2.1 |
| MT2 | 30S S5E10 | 2 | 8 | 1.0 |
| IR1 | 50S L9N11 | 2 | 8 | 4.2 |
|  | 30S S6D41 | 2 | 8 | 1.0 |
| IR2 | 50S L9N11 | 2 | 8 | 4.2 |
|  | 30S S11E76 | 2 | 8 | 1.0 |
| Control | RNaseA A9 | 3 | 4 | 10.4 |

<sup>1</sup>Percent enrichment was measured by dividing the number of mutant colonies detected by MASC-PCR and confirmed via Sanger sequencing by the total number of colonies that were screened by MASC-PCR and multiplying the result by 100.

In order to site specifically label the *E. coli* ribosome at the selected thirteen positions, we decided to genomically encode the ncAA *p*-azido-phenylalanine (*p*-AzF) using engineered UAG stop codons that are typically decoded by polypeptide release factor (RF) 1 during translation termination. The choice of *p*-AzF was driven by the fact that it undergoes a rapid and robust, bioorthogonal conjugation reaction with dibenzocyclooctyne (DBCO)-derivatized^15^ labels under mild buffer and temperature conditions (Fig. 1c). To genomically encode *p*-AzF at engineered UAG stop codons in a manner that was free of potential competition from RF1-mediated translation termination, we chose to use the C321ΔA strain of *E. coli* that has been developed by Church and co-workers^16^ (Methods). In C321ΔA, all 321 naturally occurring, essential TAG stop codons in *E. coli* have been converted to TAA stop codons that are decoded by RF2 and, moreover, the gene encoding RF1, *prfA*, has been deleted. Additionally, C321ΔA carries a temperature-inducible lambda prophage for carrying out single-stranded, DNA-guided homologous recombination (*i.e.*, lambda Red (λ_Red_)-mediated recombineering)^17^, thus allowing MGE to be used to introduce codon mutations at the desired labeling positions. To tailor C321ΔA for use with ribosomes, we disrupted the gene encoding RNase A^18–20^ (Methods), and to tailor it for *p*-AzF incorporation, we transformed it with a plasmid expressing an orthogonal nonsense suppressor tRNA that recognizes UAG codons and a tRNA synthetase specific for amino-acylating this tRNA with *p*-AzF^21, 22^ (Methods).

Starting with our tailored C321ΔA strain, we performed six to eight rounds of λ_Red_-mediated MGE using a multiplexed pool of oligonucleotides targeting the thirteen labeling sites for mutation to TAG stop codons in the ribosomal protein genes in the *E. coli* genome (Supplementary Table 1) (Methods). All rounds of MGE were performed in the presence of 1 mM *p*-AzF such that incorporation of the ncAA into the MBC occurs immediately upon introduction of the mutation into the genome. The resulting cell population was screened using multiplexed, allele-specific colony (MASC)-PCR^23^ (Methods) such that we could identify strains carrying the mutations as well as the position and nature of the mutations (Supplementary Fig. 2). Once identified, the presence of mutations was further confirmed by Sanger sequencing (Supplementary Fig. 3). Using this approach, we were able to isolate ten strains in which each strain carried either one or two TAG mutations at locations corresponding to ten of the thirteen positions we originally targeted. We hypothesized that strains carrying TAG mutations at the remaining three positions could not be isolated because these mutations conferred significant fitness disadvantages. Supporting this hypothesis, the percent enrichment of mutation at each of the targeted positions ranged between two- to ten-fold lower than a mutation of a similar size in a non-essential gene (Table 1). These observations are consistent with our expectation that, during the rounds of MGE, our approach selects for MBCs with functionally permissible mutations.

To demonstrate that the isolated, mutant *E. coli* strains were able to assemble ribosomes with site-specific incorporation of a *p*-AzF that can be efficiently labeled, we first purified ribosomes from three double-mutant strains (HS1, HS2, and IR1), and one single-mutant strain (MT1) (Methods). To ensure high-efficiency labeling, we separated the purified ribosomes into 30S and 50S subunits. The subunits were then labeled with DBCO-derivatized Cy3 and/or Cy5 fluorophores (Methods). The ribosomal proteins from each subunit were separated on an SDS-PAGE gel, and the gel was imaged using a fluorescence gel scanner (Fig. 2a). In these scans, fluorescence was observed exclusively from bands corresponding to ribosomal protein(s) whose gene(s) contained the TAG mutation(s). Moreover, fluorescence scanning of an SDS-PAGE gel containing ribosomal proteins obtained from the 30S and 50S subunits of a C321ΔA strain that did not contain any TAG mutations, but that was otherwise grown in the presence of *p*-AzF and used to purify and fluorophore-label the 30S and 50S subunits in a manner identical to that of the strains containing TAG mutations did not exhibit any fluorescence. This observation demonstrates the site-specific nature of *p*-AzF incorporation into the ribosomal proteins of the mutant strains (Supplementary Fig. 4). To measure the labeling efficiency of each labeled position, we quantified the fluorescence intensity of each labeled protein band, and compared it to a standard curve generated using known quantities of a fluorophore-labeled protein standard (Supplementary Fig. 5) (Methods). We found that the labeling efficiency was different for each targeted position, and ranged from a high of 96 % for labeling at the aspartic acid at residue position 41 on the 30S subunit protein S6 (S6 D41) to a low of 15 % for S7 G112 (Supplementary Table 2). While it’s possible that the lower labeling efficiencies could be due to low *p*-AzF incorporation caused by background suppression of the UAG mutation by near-cognate aa-tRNAs (*e.g.*, Tyr-tRNA), such background suppression has been found to be quite low in C321ΔA-based strains in presence of *p*-AzF^21^ and we therefore suspect it is more likely due to incomplete fluorophore labeling arising from the potentially low solvent exposure of the targeted residue. Regardless of the origin of any low labeling efficiency, because our approach uses MGE, it is easy to screen for another, comparable labeling position with higher labeling efficiency.

**Figure 2.**
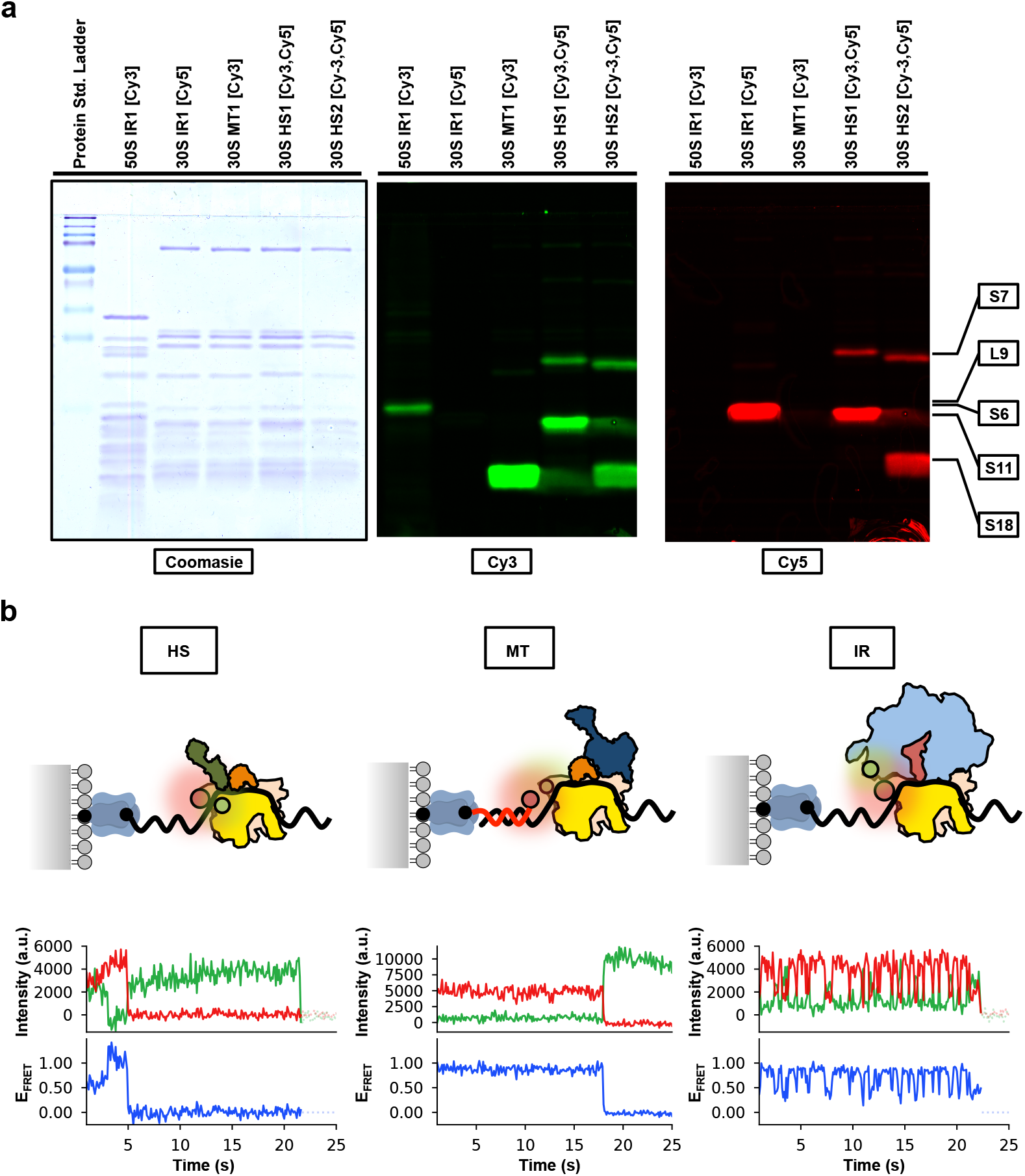
Site-specific Cy3- and/or Cy5 labeling of ribosomes purified from genomic mutant strains and smFRET experiments of ribosomal complexes assembled using labeled ribosomes. (a) SDS-PAGE analysis of ribosomal proteins derived from 30S or 50S subunits isolated from the IR1, MT1, HS1, and HS2 mutant strains and reacted with DBCO-derivatized Cy3- and/or Cy5 fluorophores. Left-hand panel shows visible light scan of Coomasie-stained gel. Middle and right-hand panels show fluorescence emission scans of pre-Coomassie-stained gel using excitation wavelengths of 532 nm for Cy3 (middle panel) and 635 nm for Cy5 (right-hand panel). The position at which each labeled ribosomal protein is expected to run on the SDS-PAGE gel was determined using a standard protein molecular weight ladder, and is indicated on the right-hand side of the figure. (b) Schematics (top row), representative Cy3- and Cy5 fluorescence intensity *versus* time trajectories (center row), and corresponding *E*_FRET_ *versus* time trajectories (bottom row) for smFRET experiments performed on ribosomal complexes assembled using Cy3- and/or Cy5-labeled 30S and/or 50S subunits isolated from the HS1 (left column), MT1 (middle column), and IR1 (right column) mutant strains. In the schematics, the surface of the microfluidic flow-cells are shown as a grey surface, PEGs are shown as grey spheres, biotinylated-PEGs are shown as black spheres, streptavidin is shown in blue-grey, mRNAs are shown as black curves, DNAs are shown as red curves, biotins at the 5’ end of the mRNAs or the 3’ end of the DNAs are shown as black spheres, the head domain of the 30S subunits is shown in yellow, the body domains of the 30S subunits are shown in tan, the 50S subunits are shown in light blue, the deacylated tRNA^fMet^ is shown in dark pink, IF1 is shown in orange, IF2 is shown in dark blue, IF3 is shown in dark green, the Cy3 fluorophores are shown as light green spheres, and the Cy5 fluorophores are shown as light red spheres. In the Cy3 and Cy5 fluorescence intensity *versus* time trajectories, the Cy3 and Cy5 fluorescence intensities are shown as green and red curves, respectively. In the *E*_FRET_ *versus* time trajectories, the *E*_FRET_ is shown as blue curves.

To demonstrate the utility of our labeled ribosomes for smFRET studies, we performed smFRET experiments using wide-field, total internal reflection fluorescence (TIRF) microscopy (Methods). In these experiments, ribosomal complexes assembled on a 5’-biotinylated mRNA using the Cy3 FRET donor- and/or Cy5 FRET acceptor-labeled ribosomes described above and any required tRNAs and/or translation factors were tethered to the surface of a polyethylene glycol (PEG)/biotin-PEG-derivatized quartz microfluidic flow-cell using a biotin-streptavidin-biotin bridge (Fig. 2b). As previously described, tethered complexes were illuminated using the evanescent wave generated by the 532 nm laser component of the TIRF microscope, and Cy3 and Cy5 fluorescence emissions from individual ribosomes were collected, wavelength separated, and imaged using the optical and detector components of the TIRF microscope^24, 25^. We imaged complexes harboring the HS1, MT1, and IR1 smFRET signals and, in each case, we observed Cy3- and Cy5 florescence intensity *versus* time trajectories that exhibited anti-correlated Cy3 and Cy5 fluorescence intensity changes followed by single-step photobleaching of the Cy3- and/or Cy5 fluorophore, demonstrating that the complexes contained single Cy3- and Cy5 fluorophores that were positioned so as to generate detectable FRET efficiencies (*E*_FRET_s) (Fig. 2c). For each complex, the *E*_FRET_ *versus* time trajectories matched our expectations regarding the *E*_FRET_ values we observed and whether the trajectories remained at a single *E*_FRET_ value or fluctuated between multiple *E*_FRET_ values as a function of time. Thus, the new smFRET signals reported here can now be used for smFRET studies of ribosomal complex dynamics that have heretofore remained difficult or impossible to interpret or perform due to the lack of a facile, versatile, and robust method for generating *in vivo* assembled, fully functional, site-specifically labeled ribosomes.

In conclusion, we have demonstrated how a combination of MGE, ncAA mutagenesis technology, and bioorthogonal chemistry can be used to label MBCs with biochemical or biophysical probes without significantly perturbing the genomic regulation, *in vivo* assembly, structure, or function of the MBCs. Our method is general, facile, versatile, and robust and can therefore be used to site-specifically label any MBC at virtually any position with almost any type of probe, enabling a wide array of biochemical and biophysical studies of MBCs. Moreover, given recent advances in the use of MGE and ncAAs in other bacterial^26^, yeast^27^, and mammalian cells^28^, our approach can be readily expanded for site-specific labeling of both bacterial and eukaryotic MBCs. In addition, combining MGE with co-selection or counter-selection methods^29^ should further improve the ease, efficiency, and success of our approach. Likewise, emerging improvements in ncAA mutagenesis technology to recode sense codons and quadruplet codons (*i.e.*, *via* +1 frameshifting) as well innovations in novel bioorthogonal chemistries should allow future expansion of the approach described here to include multi-site-specific bioorthogonal labeling of MBCs.

## ACKNOWLEDGEMENTS

We thank Harris Wang and Jimin Park for helpful discussions regarding MGE, Bridget Huang and Haixing Li for supplying useful reagents, and Colin Kinz-Thompson and Kelvin Caban for comments on the manuscript. This work was supported by funds to R.L.G. from the National Institute of Health (R01 GM084288 and R01 GM119386).

## AUTHOR CONTRIBUTIONS

B.J.D. and R.L.G. designed the experiments. B.J.D. performed the experiments. B.J.D. and R.L.G. analyzed the results. B.J.D. and R.L.G. wrote the manuscript. Both authors agreed to the final version of the manuscript.

## COMPETING INTERESTS

The authors declare no competing financial interests.

## ONLINE METHODS

### Strains and Plasmids

The C321ΔA strain was a kind gift from George Church^16^. It is an *E. coli* MG1655 strain in which all 321 naturally occurring essential TAG stop codons have been changed to TAA stop codons and the gene encoding RF1, *prfA*, has been deleted. Its exact geneotype is: Δ(ybhB-bioAB)∷[lcI857 N(cro-ea59)∷tetR-bla] ΔprfA ΔmutS∷zeoR. To render the C321ΔA strain RNase A deficient, we performed four homologous recombination cycles using a mutagenic oligonucleotide that targeted the ninth codon of the gene encoding RNAse A, *rna*, for mutation to a TAA stop codon using a previously described method^9, 23^. Strains containing the analogous mutation in *rna* are standard tools in biochemical and biophysical studies of ribosomes, translation, and translational control^18–20^. The pEvol-pAzFRS.2.t1 plasmid encoding an orthogonal nonsense suppressor tRNA that recognizes UAG codons and the tRNA synthetase that specifically amino-acylates this tRNA with *p*-AzF was a kind gift from Farren Isaacs.^21^ For the experiments described here, we transformed the RNase A-deficient C321ΔA strain with pEvol-pAzFRS.2.t1.

### Multiplexed genome engineering cycles

Thirteen oligonucleotides targeting different codon sites in ribosomal protein genes for mutation to TAG codons were designed as previously described^9, 23^, and purchased from Integrated DNA Technologies Inc. (Supplementary Table 1). MGE cycles were performed using the RNase A-deficient C321ΔA strain that had been transformed with the pEvol-pAzFRS.2.t1 plasmid and following a previously published procedure^9, 23^. All of the media used for MGE contained 1 mM *p*-AzF and 0.2 % arabinose. Cultures at the end of 6-8 MGE cycles were used to screen for mutant strains using MASC-PCR (*infra vide*). If a double mutant was needed, as in the case of generating the HS signals, one of the successfully engineered single-mutant strains was subjected to single-plexed genome engineering using the oligonucleotide targeting the second successfully engineered position.

### Screening for mutants and calculation of percent enrichment

MASC-PCR was used to screen for mutant strains in the pool of cells present in the culture after 6-8 MGE cycles, using previously published methods^9, 23^. Colonies that tested positive for mutations in the MASC-PCR screening were sequenced by Sanger sequencing (Genewiz) in order to confirm the presence and identity of the mutation. The percent enrichment of a particular mutation was calculated by dividing the number of confirmed mutant colonies by the total number of colonies screened and multiplying the result by 100.

### Ribosome purification and labeling

70S ribosomes were purified using previously established procedures^30^ with slight modifications. 1 mL culture of C321ΔA wt or isolated mutant strain was used to inoculate 1L of 2× Yeast Tryptone media (2× YT) containing 1mM *p*-AzF, and 0.2 % arabinose. The culture was grown to mid-log phase (OD 0.5), and the cells were harvested by centrifugation at 5000×g for 10 min. The actively translating 70S ribosomes and polysomes were then purified from the harvested cells as previously described^30^.

In order to efficiently label the ribosomal subunits with DBCO-derivatized Cy3 and/or Cy5, the purified 70S ribosomal pellet was gently resuspended in Low Magnesium Buffer (20 Tris(hydroxymethyl)aminomethane (Tris) acetate (OAc) at pH_RT_ = 7.5, 60 mM ammonium chloride (NH_4_Cl), 1 mM magnesium acetate (Mg(OAc)_2_), 0.5 mM ethylenediaminetetraacetic acid (EDTA), and 6 mM 2-mercaptoethanol (β-ME)) such that the 70S ribosomes would dissociate into their component 30S and 50S subunits and the resulting solution was diluted such that the final concentrations of 30S and 50S subunits was 1 μM each. DBCO-derivatized Sulfo-Cy3 and/or DBCO-derivatized Sulfo-Cy5 (Lumiprobe) were then added to the diluted 30S and 50S subunits to a final concentration of 10 μM. The labeling reaction was carried out at 4 °C for 12 hours in the dark. The labeling reaction was then dialyzed against Low Magnesium Buffer to remove excess, unconjugated fluorophores. Labeled 30S and/or 50S subunits were then purified from the labeling reactions as previously reported using sucrose gradient ultracentrifugation in Low Magnesium Buffer^25^. Labeling efficiencies were calculated by interpolating the cumulative fluorescence intensity of each labeled protein band from an SDS-PAGE gel of labeled 30S and/or 50S subunits from linear standard curves generated using Cy3- or Cy5-labeled protein standards of defined quantities and labeling efficiencies. Specifically, the Cy3- and Cy5-labled protein standards we used were L1 Q18C-Cy3 (L1 [Cy3]) and RF1 S167C-Cy5 (RF1 [Cy5]), respectively, that were isolated, labeled, and purified so as to exhibit 100 % Cy3- or Cy5 labeling efficiencies as previously described^24, 31^ (Supplementary Fig. 4).

### Preparation of mRNAs, tRNAs, and translation factors

Ribosomal complexes for smFRET experiments using the HS and IR smFRET signals were assembled on a previously described^32, 33^, 5’-biotinylated model mRNA that is a variant of the mRNA encoding bacteriophage T4 gene product 32 (Bio-mRNA). Bio-mRNA was chemically synthesized (IDT) (Supplementary Table 3). Ribosomal complexes for smFRET experiments using the MT smFRET signal were assembled on a previously described^30^, non-biotinylated model mRNA consisting of the first twenty codons of bacteriophage T4 gene product 32 (NonBio-mRNA) that was hybridized to a previously described^30^, 5’-Cy5-labeled, 3’-biotinylated DNA oligonucleotide (Cy5-DNA-Bio) (Supplementary Table 3). NonBio-mRNA was synthesized by *in vitro* transcription using bacteriophage T7 RNA polymerase, as previously described^25^, and Cy5-DNA-Bio was chemically synthesized (IDT). Hybridization of NonBio-mRNA with Cy5-DNA-Bio was performed as previously described^25^. The hybridized NonBio-mRNA:Cy5-DNA-Bio was purified away from excess, unhybridized Cy5-DNA-Bio using size-exclusion chromatography. *E.coli* initiator, formylmethionine-specific tRNA (tRNA^fMet^) was purchased from MP Bio and was aminoacylated and formylated using methionyl-tRNA synthetase and methionyl-tRNA formyltranferase, respectively, using previously published procedures^30^. *E. coli* translation initiation factors (IFs) 1, 2, and 3 were expressed and purified using established protocols as described elsewhere^25^.

### Preparation of ribosomal complexes for smFRET experiments

To perform smFRET experiments using the HS smFRET signal, we attempted to prepare a 30S initiation complex (IC) similar to one that has been described as exhibiting HS dynamics in a recent cryo-EM study by Ramakrishnan and co-workers^34^. To do this, we adapted general protocols that we have previously developed for the preparation of 30S ICs^35, 36^. Specifically, we incubated 0.6 μM 30S subunits labeled with Cy3 and Cy5 (30S HS1 [Cy3/Cy5]), 1.8 μM Bio-mRNA (Supplementary Table 3), 0.9 μM fMet-tRNA^fMet^, 0.9 μM IF1, and 0.6 μM IF3 in Tris-Polymix Buffer (50 mM Tris-OAc at pH_RT_ = 7.5, 100 mM KCl, 5 mM ammonium acetate (NH_4_OAc), 5 mM Mg(OAc)_2_, 0.1 mM EDTA, 1 mM guanosine triphosphate (GTP), 5 mM putrescine-HCl, 1 mM spermidine-free base, and 6 mM β-ME) for 10 minutes at 37 °C and subsequently transferred the reaction to ice for an additional 5 minutes. The complexes were divided into 1 μL aliquots and flash-frozen by immersing the tube in liquid nitrogen. The aliquots were stored at –80 °C for future use^35, 36^. Given these reaction conditions, the fact that IF1 and IF3 were maintained at 1 μM and 25 nM, respectively, in the buffers used throughout all complex formation, dilution, tethering, and imaging steps (*vide infra*), and the results of previous biochemical and smFRET work by us and others^32^, we expect this reaction to predominantly yield the IF1- and IF3-containing 30S IC that is schematized on the left-hand panel of Figure 2b.

To perform smFRET experiments using the MT smFRET signal, we attempted to prepare a 30S IC similar to one that has been described in a recent smFRET study by Gonzalez and co-workers^32^, that would serve as a preliminary proof-of-concept model for MT signal. To do this, we again adapted general protocols that we have previously developed for the preparation of 30S ICs^32, 35, 36^. Specifically, we incubated 0.6 μM 30S subunits labeled with Cy3 (30S MT1 [Cy3]), 1.8 μM NonBio-mRNA:Cy5-DNA-Bio (Supplementary Table 3), 0.9 μM fMet-tRNA^fMet^, 0.9 μM IF1, and 0.9 μM IF2 in Tris-Polymix Buffer for 10 minutes at 37 °C and subsequently transferred the reaction to ice for an additional 5 minutes. The complexes were divided into 1 μL aliquots and flash-frozen by immersing the tubes in liquid nitrogen. The aliquots were stored at –80 °C for future use^35, 36^. Given these reaction conditions, the fact that IF1 and IF2 were maintained at 1 μM and 25 nM, respectively, in the buffers used throughout all complex formation, dilution, tethering, and imaging steps, and the results of previous smFRET studies^32^, we expect this reaction to predominantly yield the IF1-, IF2-, and fMet-tRNA^fMet^-containing 30S IC that is schematized on the center panel of Figure 2b.

To perform smFRET experiments using the IR smFRET signal, we attempted to prepare a 70S pre-translocation complex mimic lacking a peptidyl-tRNA in the ribosomal aminoacyl-tRNA binding site (*i.e.*, a 70S PRE^−A^ complex) similar to one that has been described as exhibiting IR dynamics in a previous smFRET study by Ha and co-workers^12^. To do this, we adapted general protocols that we have previously developed for the non-enzymatic preparation of 70S PRE^−A^ complexes^33^. Specifically, we incubated 15 pmol 30S subunits labeled with Cy5 (30S IR1 [Cy5]), 30 pmol Bio-mRNA (Supplementary Table 3), and 20 pmol deacylated tRNA^fMet^ in 30 μL of 70S PRE^−A^ Assembly Buffer (50 mM Tris hydrochloride (Tris-HCl) at pH_RT_ = 7.5, 70 mM NH_4_OAc, 30 mM KCl, 6 mM β-ME, and 7 mM MgCl_2_) for 10 minutes at 37 °C, at which point 10 pmol 50S subunits labeled with Cy3 (50S IR1 [Cy3]) were added and the reaction incubated for additional 20 minutes at 37 °C. The reaction was then placed on ice for 5 minutes and diluted to a final volume of 100 μL with Tris-Polymix Buffer that had been adjusted to 20 mM Mg(OAc)_2_. The reaction was carefully layered on top of 10-40% (w/v) sucrose gradient made in Tris-Polymix Buffer adjusted to 20 mM Mg(OAc)_2_ and purified by density gradient ultracentrifugation as previously described^24, 33^. The complexes were divided into 25 μL aliquots and flash-frozen by immersing the tubes in liquid nitrogen. The aliquots were stored at –80 °C for future use. Given these reaction conditions and the results of previous smFRET studies^33^, we expect this reaction to predominantly yield the 70S PRE^−A^ that is schematized on the right-hand panel of Figure 2b.

### TIRF-based smFRET experiments

Ribosomal complexes were diluted to ~100-500 pM in Tris-Polymix Buffer adjusted to 5 mM Mg(OAc)_2_ for smFRET experiments using the HS1 and MT1 smFRET signals and to 15 mM Mg(OAc)_2_ for smFRET experiments using the IR1 smFRET signal and supplemented with 1 μM IF1 and 25 nM IF3 for smFRET experiments using the HS1 smFRET signal and 1 μM IF1 and 25 nM IF2 for smFRET experiments using the MT1 smFRET signal. The complexes were tethered to the PEG/Biotin-PEG-derivatized surfaces of our quartz microfluidic flow-cells *via* a biotin-streptavidin-biotin bridge by incubating the diluted complexes in our flow-cells for 5 minutes, after which unbound complexes were flushed out of the flow-cells using the same Tris-Polymix Buffers that were used to dilute the complexes, but that had been further supplemented with an oxygen-scavenging system (5 mM protocatechuic acid and 10 nM protocatechuate-3,4-dioxygenase) and a triplet state quencher (1 mM cyclooctatetraene).

The tethered complexes were then imaged using a laboratory-built, wide-field, prism-based TIRF microscope with a diode-pumped solid-state 532 nM laser (Laser Quantum GEM532) as an excitation source for Cy3. Fluorescence emissions from Cy3 and Cy5 were collected through a 60× magnification, water-immersion objective with a numerical aperture of 1.2 (Nikon), wavelength separated using a Dual-View image-splitting device (Photometrics), and imaged using an water-cooled, electron-multiplying charged coupled device (EMCCD) camera (Andor iXon Ultra 888) operating with 2× binning. 600-frame movies were collected at a time resolution of 0.1 second using μManager^37^.

TIRF movies were analyzed using custom-written software (manuscript in preparation; Jason Hon, Colin Kinz-Thompson, R.L.G.). First, fluorophores were identified by locating local maxima pixels in the movie and classifying them into either ‘fluorophore’ or ‘background’ classes. The Cy3 and Cy5 imaging channels were then aligned by applying a polynomial transformation that had been separately computed using a control image of fiducial markers. Using the aligned Cy3 and Cy5 imaging channels, we next fit each Cy3 and Cy5 fluorophore in each image of the movie to a 2D Gaussian in order to estimate the Cy3 and Cy5 fluorescence intensity *versus* time trajectories for each identified and aligned pair of Cy3 and Cy5 fluorophores. For each time point, Cy5. fluorescence intensity values were corrected for Cy3 bleedthrough by subtracting 5% of the Cy3 florescence intensity value in the corresponding Cy3 fluorescence intensity trajectory. *E*_FRET_ *versus* time trajectories were then generated by using the Cy3 fluorescence intensity trajectories and bleedthrough-corrected Cy5 fluorescence intensity trajectories to calculate the *E*_FRET_ value at each time point of the corresponding *E*_FRET_ trajectory. The *E*_FRET_ values were calculated by dividing the Cy5 fluorescence intensity (*I*_Cy5_) by the sum of the Cy3 and Cy5 fluorescence intensities (*I*_Cy5_ + *I*_Cy3_), as previously described^38^. Visual inspection was then used to select only those *E*_FRET_ trajectories for which the corresponding Cy3 and Cy5 fluorescence intensity trajectories exhibited single-step photobleaching of both the Cy3 and Cy5 fluorophores, thereby confirming that the *E*_FRET_ trajectory originated from a pair of single donor and acceptor fluorophores.

## Data Availability

The data supporting the findings of this study are available from the corresponding author upon request.

## Code Availability

The code used to analyze the TIRF movies in this study is available from the corresponding author upon request.

