## Supplementary Information for "Multiplexed, bioorthogonal labeling of multicomponent, biomolecular complexes using genomically encoded, non-canonical amino acids"

**Supplementary Figures**

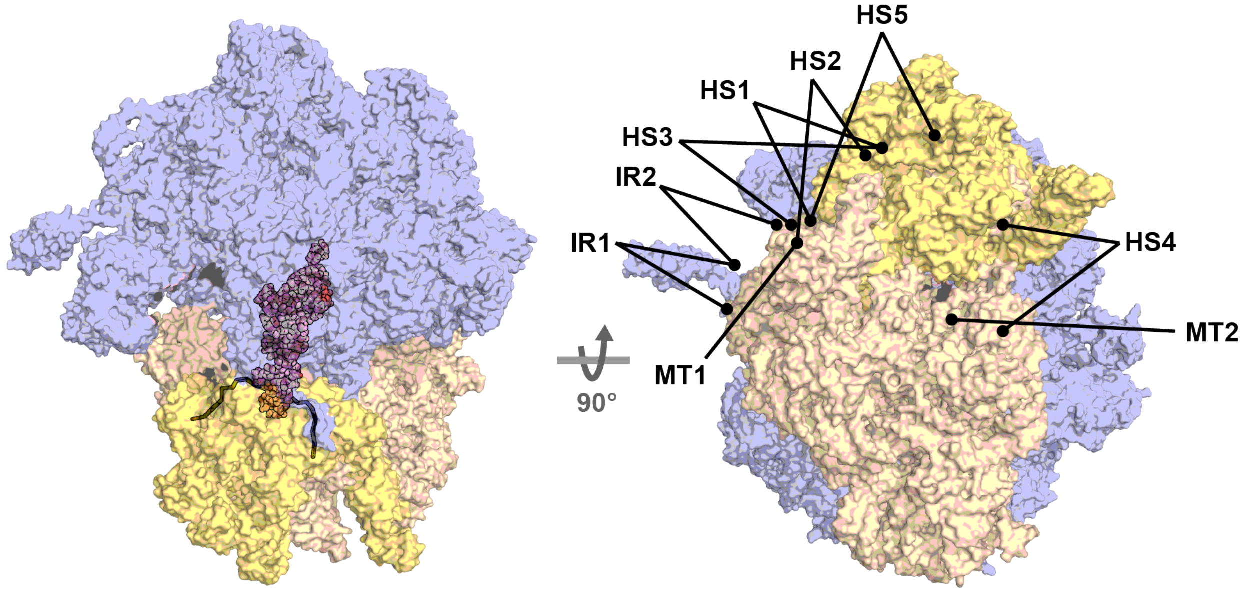


**Supplementary Figure 1. Locations of ribosomal protein residues that were targeted for labeling with FRET donor and/or acceptor fluorophores for smFRET experiments**. Target residues were selected by assessing: (*i*) their ability to report on the ribosome dynamics of interest as determined by comparative analyses of the available ribosomal complex structures, (*ii*) their relatively low phylogenetic conservation among bacterial species most closely related to *E. coli*, (*iii*) their seeming lack of participation in protein structural elements that could possibly be critical for maintaining the integrity of the targeted ribosomal protein, and (*iv*) their presumed accessibility to solvent and small-molecule, chemical labeling reagents. The targeted residues in the figure are denoted by the black spheres at the ends of the black lines and are labeled according to the ribosome dynamics that they are expected to report on: intra-subunit rotation of the head domain of the 30S subunit relative to the body domain (*i.e.*, 'head swiveling', HS); movement of a translating ribosome along its mRNA template (*i.e.*, 'mRNA translocation', MT); and rotation of the 50S subunit relative to the 30S subunit (*i.e.*, 'intersubunit rotation', IR). The structure shown here is that of an atomic-resolution, X-ray crystallographic structure of a *Thermus thermophilus* ribosomal complex (PDB ID: 5IBB) that is shown as a space-filling, surface contour model. The head domain of the 30S subunit is shown in yellow, the body domain of the 30S subunit body domain is shown in tan, the 50S subunit is shown in cyan, and the P site-bound tRNA is shown in red.

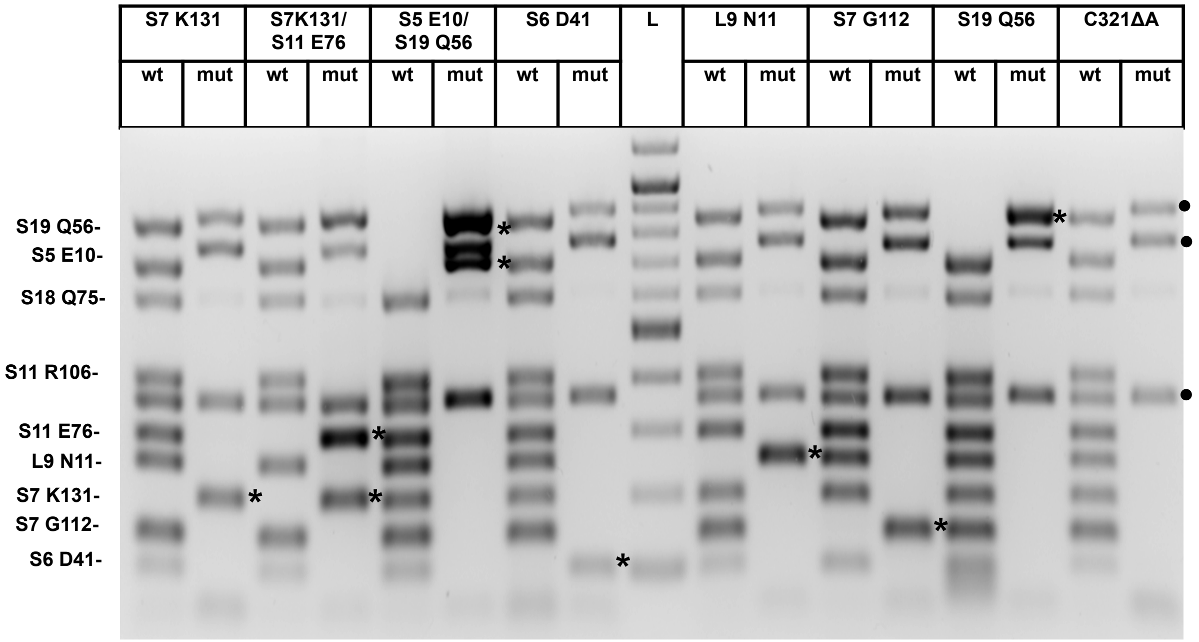


**Supplementary Figure 2. MASC-PCR-based identification of colonies containing genomic TAG mutations in ribosomal protein genes isolated after six to eight rounds of MGE.** 1.5 % agarose gel of MASC-PCR reactions performed using primers targeting either wild-type (wt) or mutant (mut) alleles and isolated single bacterial colonies as templates. Primers were designed such that each colony carrying a TAG mutation would give a uniquely sized PCR product (as indicated on the left-hand side of the gel). In these MASC-PCR reactions, the presence of a particular TAG mutation is indicated by the presence of a PCR product of a size corresponding to the TAG mutation in the lane labeled mut (indicated by the *s) and the concomitant absence of a PCR product of the same size in the lane labeled wt. Extraneous PCR products that were present in all wt or mut reactions were identified by performing MASC-PCR reactions on our tailored, parent C321ΔA strain (indicated by the •s).

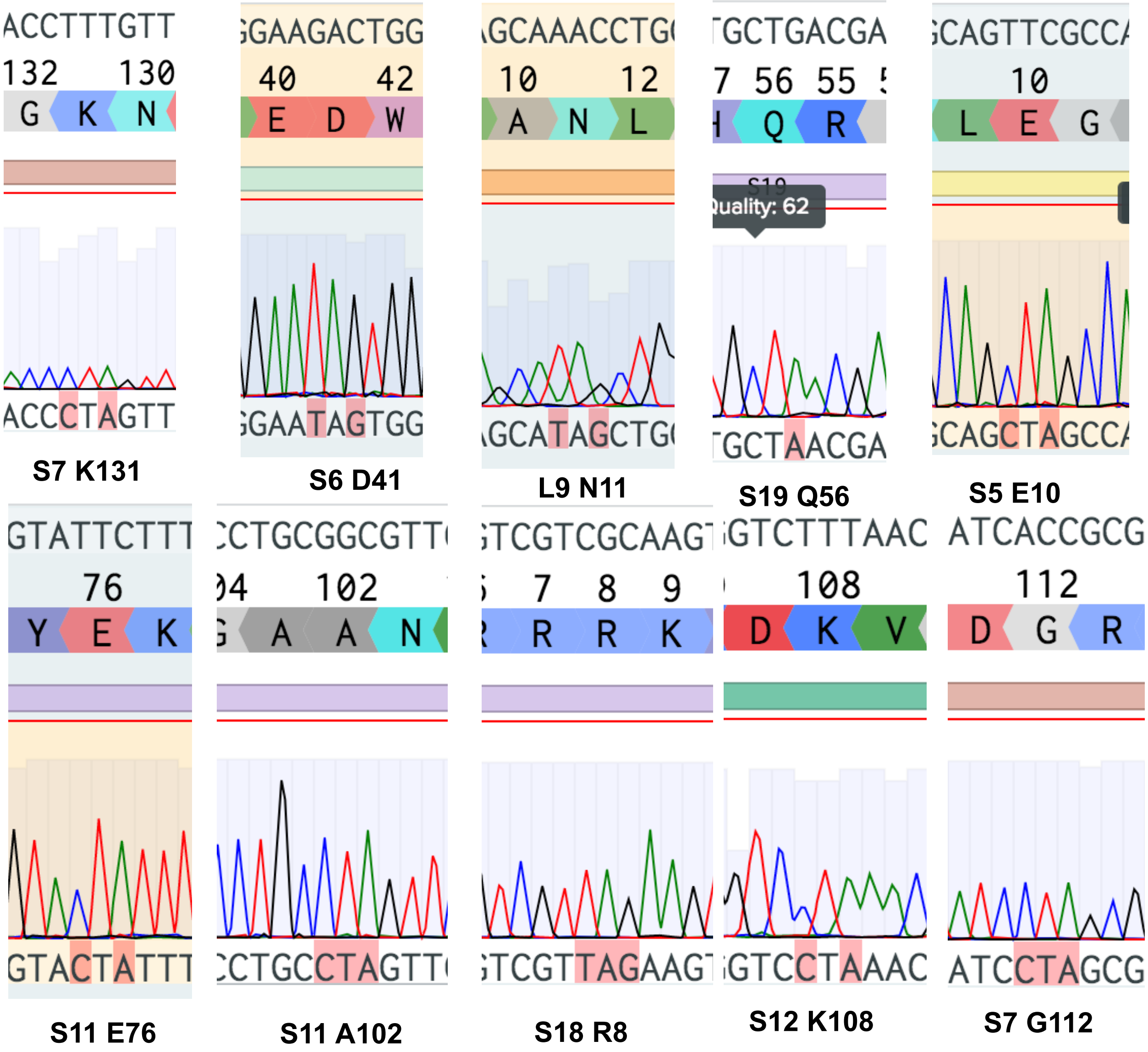


**Supplementary Figure 3. Sanger sequencing-based confirmation of colonies containing genomic TAG mutations in ribosomal protein genes isolated after six to eight rounds of MGE.** Each panel shows an alignment of the wildtype reference nucleotide- and amino acid sequences of the targeted ribosomal protein gene (top two rows), Sanger sequencing trace (middle row), and nucleotide sequence corresponding to the Sanger sequencing trace (bottom row) for PCR products amplified from the genomic mutation region using isolated mutant colonies initially identified in the MASC-PCR screening. The amino acid sequences are shown as single letter codes. The Sanger sequencing traces are color-coded based on nucleotide: A is shown in green, T is shown in red, C is shown in blue, and G is shown in black. The successfully mutated nucleotides are highlighted in pink in the nucleotide sequence corresponding to the Sanger sequencing trace.

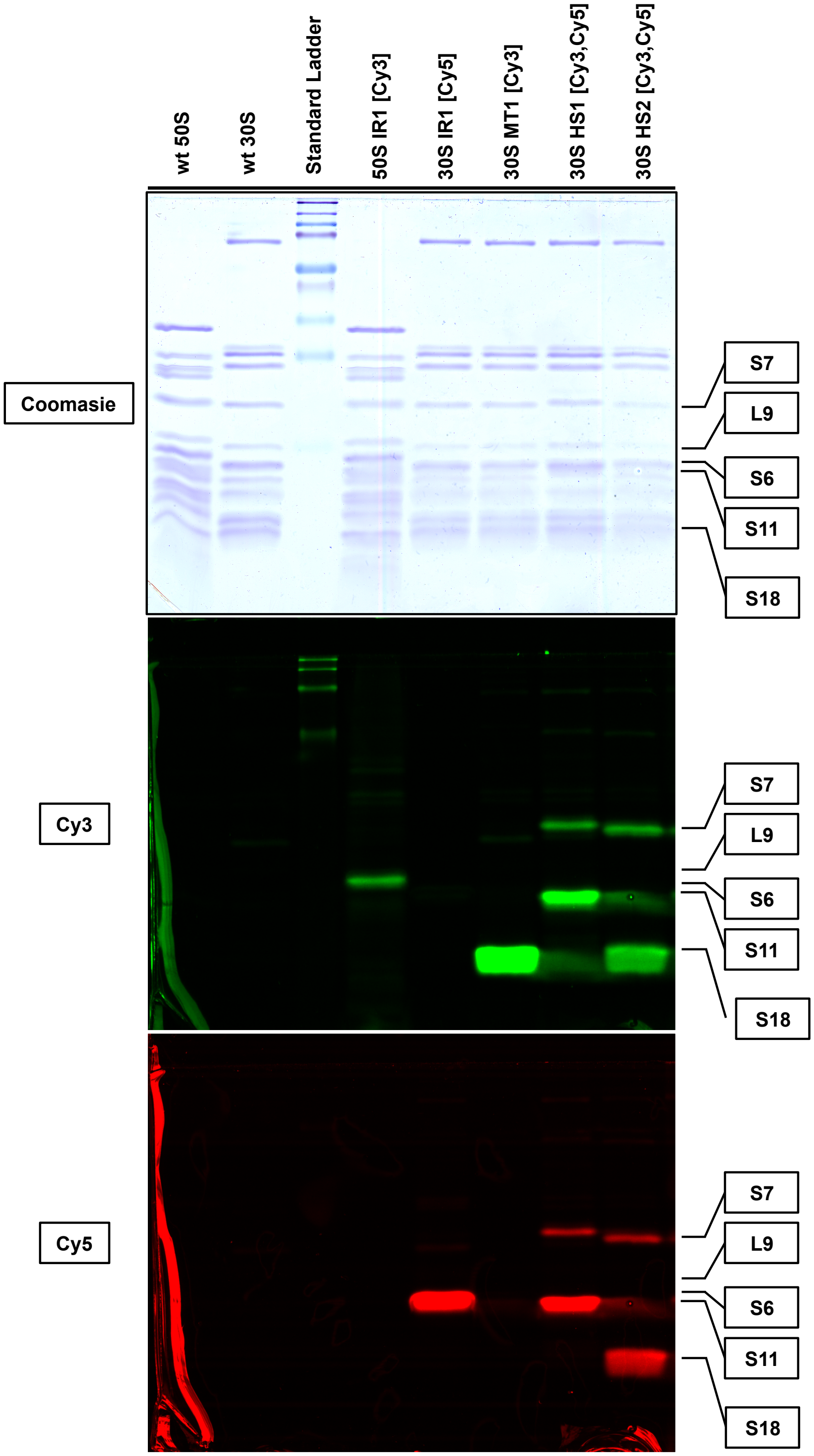


**Supplementary Figure 4. SDS-PAGE analysis of ribosomal proteins derived from 30S or 50S subunits isolated from our tailored, parent C321ΔA strain (wt) and the IR1, MT1, HS1,**

**and HS2 mutant strains and reacted with DBCO-derivatized Cy3- and/or Cy5 fluorophores.** Top panel shows visible light scan of Coomasie-stained gel. Middle and bottom panels show fluorescence emission scans of pre-Coomassie-stained gel using excitation wavelengths of 532 nm for Cy3 (middle panel) and 635 nm for Cy5 (bottom panel). The position at which each labeled ribosomal protein is expected to run on the SDS-PAGE gel was determined using a standard protein molecular weight ladder (Lane 3) and is indicated on the right-hand side of the figure.

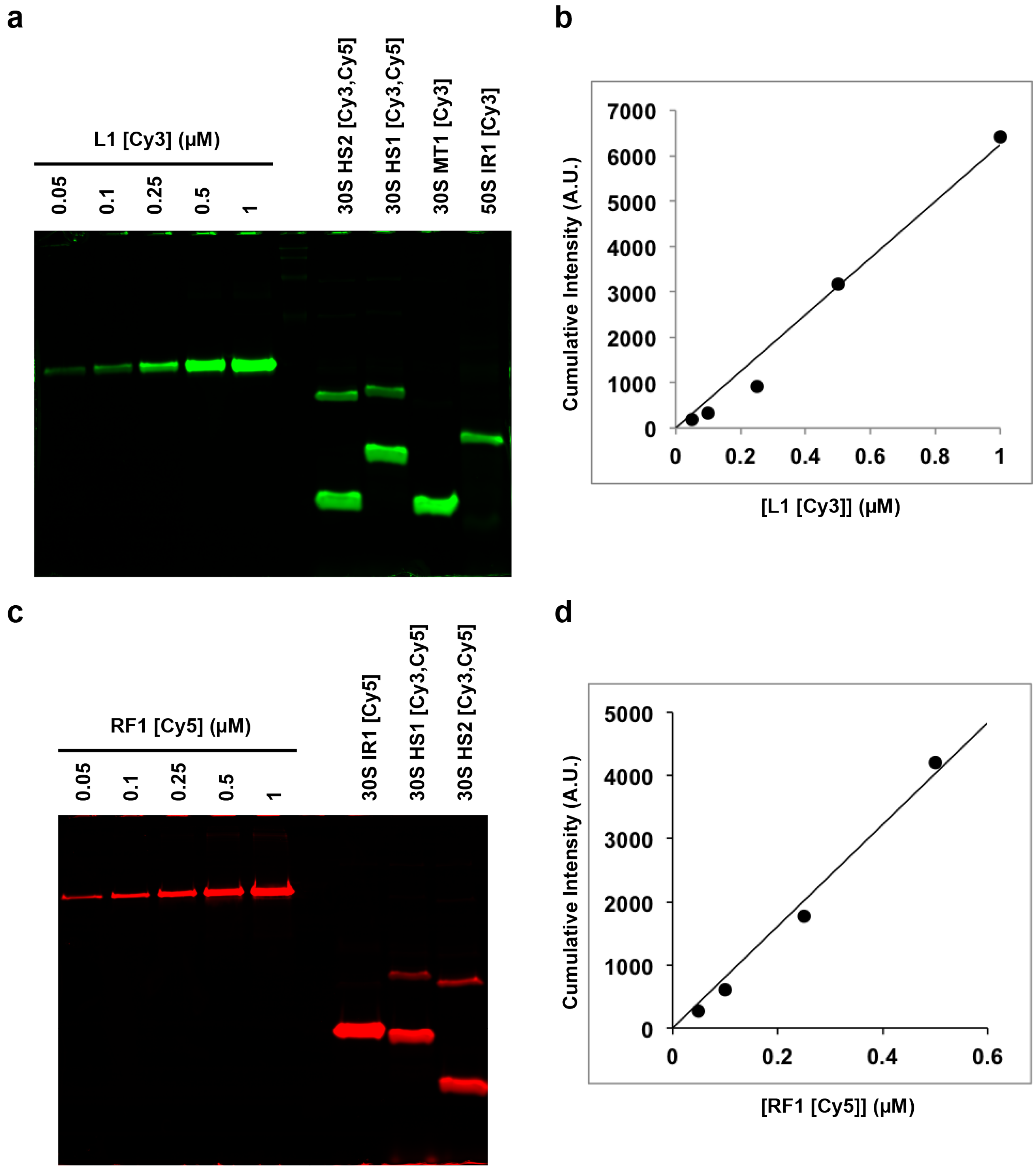


**Supplementary Figure 5. Quantification of labeling efficiency for 30S and 50S subunits purified from HS1, HS2, MT1, and IR1 mutant strains.** Fluorescence emissions scans at excitation wavelengths of 532 nm for (a) Cy3 and 635 nm for (c) Cy5 of SDS-PAGE gels containing 12 μl of the indicated concentrations of Cy3- or Cy5-labeled protein standards (L1

[Cy3] and RF1 [Cy5], respectively) and 12 µl of 1 µM of Cy3- and/or Cy5-labeled 30S or 50S subunits isolated from the indicated mutant strains as described in Methods. Linear standard curves plotted using the cumulative fluorescence intensities of the protein bands corresponding to different concentrations of (b) L1 [Cy3] and (d) RF1 [Cy5]. The data points (black filled circles) in each standard curve were fit to a linear function of the form y = mx + b (black lines), where y is the cumulative fluorescence intensity, x is the concentration of L1 [Cy3] or RF1 [Cy5], m is the slope, and b is y-intercept, which was required to go through zero. The equation for the standard curve describing L1 [Cy3] was y = 6241x + 0 and fit the data with an $R^2$ of 0.98 and the equation for the standard curve describing RF1 [Cy5] was y = 8063.5x + 0 and fit the data with an $R^2$ of 0.98.

### Supplementary Tables

**Supplementary Table 1.** The DNA oligonucleotides used for MGE. The nucleotides corresponding to the mutation are denoted in red and phosphorothioated nucleotides are followed by an asterisk (*).

| Target Residue | DNA Oligonucleotide Sequence |
|---|---|
| **S7 G112** | G*C*A*T*CAGAAAGTTCGTTCGCCAGGCGCAGAGCCATGGATTTATC**CTA**GCGTTTACGAGCAGCTTCAACGATCCAACGCATTGCCAGAGCAT |
| **S7 K131** | A*C*G*T*TTCTTAACTGCAGTACC**C**T**A**GTTTTCTGCAGCATCAGAAAGTTCGTTCGCCAGGCGCAGAGCCATGGATTTATCACCGCGTTTACGA |
| **S11 A102** | A*A*T*G*TTAGTGATGCGGAAACCTGC**CTA**GTTCAGAGCACGAATAGTAGATTCGCGGCCTGGACCCGGACCTTTAACCATAACTTCCAGATT |
| **S18 Q75** | T*A*C*C*TTATCCTCTCAAAGTCGTATTAATGGACCGTGACCGATTACT**A**ATGGCGATCAGTGTACGGCAGCAGGGACAGGTAGCGAGCGCGTT |
| **S18 R8** | T*A*G*T*CGATCTCTTGAACGCCTTCCGCGGTGAAACGGCAGAACTT**CTA**ACGACGGAAATAACGTGCCATATGGTTAGTCTCCAGAATCTATC |
| **S6 D41** | T*T*G*T*GCAGTTTGTTGATCGGGTAAGCCAGCTGACGGCGGCCCCA**C**T**A**TTCCAGACGGTGGATCTTGCCTTCTGCACCAGTGATGGCAGCAG |
| **L9 N11** | G*C*A*T*AGCCCGCTTTAACGTTTACCTGATCACCCAGGCTACCCAG**C**T**A**TGCTACTTTATCAAGCAGAATAACTTGCATTACCTTATCCTCTC |
| **S11 E76** | G*G*A*C*CCGGACCTTTAACCATAACTTCCAGATTCTTGATGCCGTA**CTA**TTTCACGGCGTCAGCGCAACGCTCTGCTGCAACCTGAGCTGCAA |
| **S5 E10** | A*C*G*G*TTTTAGATACGCGGTTTACCGCGATCAGCTTTTCCTGCAG**C**T**A**GCCAGCTTGTTTTTCGATGTGAGCCATCTTACACCTCTACCTTA |
| **S12 K108** | G*A*A*C*GAGCCTGCTTACGGTC**C**T**A**AACGCCGGAGCAGTCAAGCGCACCACGTACGGTGTGGTAACGAACACCCGGGAGGTCTTTAACACGA |
| **S19 Q56** | A*G*T*T*TGTGACCAACCATTTCGTCGGTTACAAATACCGGAACGTGCT**A**ACGACCATTATGGACAGCGATGGTCAAACCGATCATGTTAGGAA |
| **S13 D11** | TGCCGACGCCATAAATCGAAGTTAATGCGATTACGGCATGCTTATG**C**T**A**AGGAATGTTAATGCCTGCTATACGGGCCACTATGCACTCCT |
| **S11 R106** | A*A*C*C*GTTATGAGGGATCGGAGTCACATCAGTAATGTTAGTGAT**CTA**GAAACCTGCGGCGTTCAGAGCACGAATAGTAGATTCGCGGCCTG |
| **RNase A A9** | G*T*A*T*GAGTTCCACACCCATTATGAAAGCATTCTGGCGTAACGCC**TAA**TTGCTCGCGGTTTCTCTGCTTCCCTTCTCTTCTGCCAACGCCT |

**Supplementary Table 2.** Labeling efficiencies of targeted positions quantified by interpolating the cumulative fluorescence intensity of each labeled protein band from an SDS-PAGE gel of labeled 30S and/or 50S subunits from a linear standard curve generated using Cy3- or Cy5-labeled protein standards of defined quantities and labeling efficiencies (Methods, Supplementary Fig. 5).

| smFRET Signal | Labeling Position | Labeling Efficiency (%) |
|---|---|---|
| **IR1** | 50S L9N11 | 30 |
| | 30S S6D41 | 96 |
| **HS1** | 30S S7G112 | 15 |
| | 30S S11A102 | 75 |
| **HS2** | 30S S7K131 | 22 |
| | 30S S18R8 | 71 |
| **MT1** | 30S S18R8 | 71 |

**Supplementary Table 3.** Sequences of mRNAs and DNA oligonucleotide used in smFRET experiments.

| Name | Sequence |
|---|---|
| Bio-mRNA | 5’-Biotin-CAACCUAAAACUUACACAAAUUAAAAAGGAAAUAGACAUGUUC AAAGUCGAAAAAUCUACUGCU-3’ |
| NonBio-mRNA | 5’-GGCAACCUAAAACUUACACAGGGCCCUAAGGAAAUAAAAAUGUUUA AACGUAAAUCUACUGCUGAACUCGCUGCACAAAUGGCUAAACUGAAU GGCAAUUAAGGAUC-3’ |
| Cy5-DNA-Bio | 5’-Cy5-TGTAAGTTTTAGGTTGCC-Biotin-3’ |
